# Bispecific antibody neutralizes circulating SARS-CoV-2 variants, prevents escape and protects mice from disease

**DOI:** 10.1101/2021.01.22.427567

**Authors:** Raoul De Gasparo, Mattia Pedotti, Luca Simonelli, Petr Nickl, Frauke Muecksch, Irene Cassaniti, Elena Percivalle, Julio C. C. Lorenzi, Federica Mazzola, Davide Magrì, Tereza Michalcikova, Jan Haviernik, Vaclav Honig, Blanka Mrazkova, Natalie Polakova, Andrea Fortova, Jolana Tureckova, Veronika Iatsiuk, Salvatore Di Girolamo, Martin Palus, Dagmar Zudova, Petr Bednar, Ivana Bukova, Filippo Bianchini, Dora Mehn, Radim Nencka, Petra Strakova, Oto Pavlis, Jan Rozman, Sabrina Gioria, Josè Camilla Sammartino, Federica Giardina, Stefano Gaiarsa, Qiang Pan Hammarström, Christopher O. Barnes, Pamela J. Bjorkman, Luigi Calzolai, Antonio Piralla, Fausto Baldanti, Michel C. Nussenzweig, Paul D. Bieniasz, Theodora Hatziioannou, Jan Prochazka, Radislav Sedlacek, Davide F. Robbiani, Daniel Ruzek, Luca Varani

**Author notes:** These authors contributed equally (Raoul De Gasparo, Mattia Pedotti).

## Abstract

Neutralizing antibodies targeting the receptor binding domain (RBD) of the SARS-CoV-2 Spike (S) are among the most promising approaches against coronavirus disease 2019 (COVID-19)^1,2^. We developed a bispecific, IgG1-like molecule (CoV-X2) based on two antibodies derived from COVID-19 convalescent donors, C121 and C135^3^. CoV-X2 simultaneously binds two independent sites on the RBD and, unlike its parental antibodies, prevents detectable S binding to Angiotensin-Converting Enzyme 2 (ACE2), the virus cellular receptor. Furthermore, CoV-X2 neutralizes SARS-CoV-2 and its variants of concern, as well as the escape mutants generated by the parental monoclonals. In a novel animal model of SARS-CoV-2 infection with lung inflammation, CoV-X2 protects mice from disease and suppresses viral escape. Thus, simultaneous targeting of non-overlapping RBD epitopes by IgG-like bispecific antibodies is feasible and effective, combining into a single molecule the advantages of antibody cocktails.

---

The COVID-19 pandemic prompted an unprecedented effort to develop effective countermeasures against SARS-CoV-2. Pre-clinical data and phase III clinical studies indicate that monoclonal antibodies (mAbs) could be effectively deployed for prevention or treatment during the viral symptoms phase of the disease^1,2^. Cocktails of two or more mAbs are preferred over a single antibody for increased efficacy and prevention of viral escape. However, this approach requires increased manufacturing costs and volumes, which are problematic at a time when the supply chain is under pressure to meet the high demand for COVID-19 therapeutics, vaccines and biologics in general^4^. Cocktails also complicate formulation^5,6^ and hinder novel strategies like antibody delivery by viral vectors or by non-vectored nucleic acids^7-9^. Instead, multispecific antibodies embody the advantages of a cocktail within a single molecule.

To this avail, we employed structural information^10^ and computational simulations to design bispecifics that would simultaneously bind to (i) independent sites on the same RBD and (ii) distinct RBDs on a S trimer. Out of several designs evaluated by atomistic Molecular Dynamics simulations, 4 were produced and CoV-X2 was the most potent neutralizer of SARS-CoV-2 pseudovirus, with half-maximal inhibitory concentration (IC_50_) = 0.04 nM (5.8 ng/mL) (Extended Data Fig.1). CoV-X2 is a human-derived, CrossMAb-format IgG1-like bispecific antibody^11^ resulting from the combination of the Fragment antigen binding (Fab) of mAbs C121 and C135, two potent SARS-CoV-2 neutralizers^3^. Structural predictions showed that CoV-X2, but not its parental monoclonals, can bind bivalently to all RBD conformations on the S trimer, preventing ACE2 access (Fig.1a and Extended Data Fig.2)^12^.

**Fig. 1.**
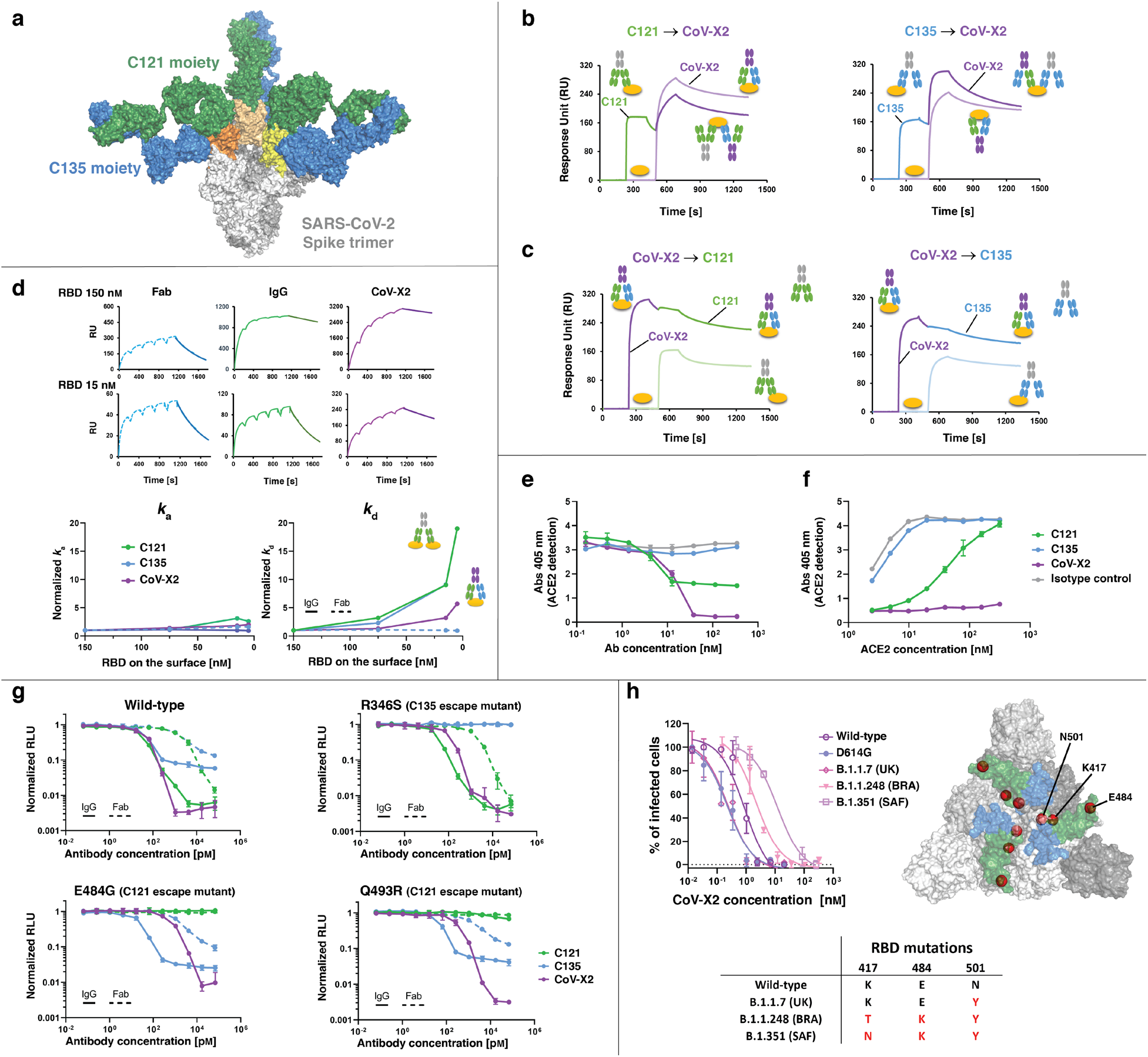
Biochemical and *in vitro* neutralizing properties of CoV-X2 are superior to its parental mAbs. **a**, Computational simulations predict bivalent binding of CoV-X2 to all three RBDs on the S trimer (see also Extended Data Fig.2). Green and blue are C121 and C135 moieties, respectively; RBDs are in shades of yellow/orange. **b, c**, SPR demonstrates that both arms of CoV-X2 are functional. In (**b**), immobilized RBD complexed with the indicated mAb (first antibody) binds to CoV-X2 (second antibody). In (**c**), the RBD/CoV-X2 complex prevents binding by the single mAbs. Shaded colors are controls (second antibody only). **d**, Both arms of CoV-X2 bind simultaneously to the RBD since, contrary to the monoclonals, avidity is retained at decreasing RBD concentrations. On top, representative SPR traces indicating the different dissociations of antibodies (or Fab) binding to RBD immobilized at different concentrations on the SPR chip (see also Extended Data Fig.6). At the bottom, plots of the normalized *k*_a_ and *k*_d_ values obtained with different concentrations of immobilized RBD. Increasing normalized dissociation rate (*k*_d_) values indicate loss of avidity. **e, f**, CoV-X2 fully prevents ACE2 binding to S trimer in ELISA. ACE2 binding to antibody/S trimer complexes is measured either with increasing concentration of the indicated antibody and constant ACE2 (**e**), or at constant antibody concentration with increasing ACE2 (**f**). Mean with standard deviation of two experiments is shown. **g**, CoV-X2 neutralizes SARS-CoV-2 pseudovirus and escape mutants of its parental mAbs. Normalized relative luminescence (RLU) for cell lysates after infection with nanoluc-expressing SARS-CoV-2 pseudovirus in the presence of increasing concentrations of antibodies. Wild-type SARS-CoV-2 pseudovirus (left) is shown alongside three escape mutants generated in the presence of C121 or C135^15^. Dashed lines are parental Fabs. Mean with standard deviation; one of two independent experiments. **h**, Neutralization of SARS-CoV-2 isolates with sequences corresponding to viruses first isolated in China (wild-type), Italy (D614G), United Kingdom (UK; B.1.1.7), Brazil (BRA; B.1.1.248) and South Africa (SAF; B.1.351). RBD residues mutated in the variants are indicated in the table and as red spheres on the S trimer structure, where the epitope of C135 (blue) and C121 (green) are shown.

CoV-X2 bound with low nanomolar affinity to RBD, S trimer, and to several mutants, including the naturally occurring variants B.1 (D614G in S protein), B.1.1.7 (N501Y in RBD) and B.1.351 (K417N, E484K and N501Y in RBD)^13,14^, and the escape mutants of the parental mAbs^15^ (Extended Data Figs.3-5).

CoV-X2 also bound to pre-formed C121/RBD and C135/RBD complexes, thus confirming that both of its arms are functional (Fig.1b,c). Next, an avidity assay by Surface Plasmon Resonance (SPR) was used to experimentally confirm the computational prediction that CoV-X2 can simultaneously engage two sites on the same RBD (Methods, Fig.1d and Extended Data Fig.6). Avidity occurs when IgGs bind bivalently to antigens, resulting in slower dissociation rates (*k*_d_) (Extended Data Fig.6a). Accordingly, C121 and C135 IgG showed avidity at high antigen concentrations due to inter-molecular binding of adjacent RBDs; at lower antigen concentrations the dissociation rate was instead faster since inter-molecular binding was prevented by the increased distance between RBD molecules, resulting in loss of avidity. Intra-molecular avidity is not possible for C121 and C135 since a single epitope is available on each RBD molecule. By contrast, CoV-X2 maintained avidity even at low antigen concentrations, indicating bivalent, intra-molecular binding (Fig.1d and Extended Data Fig.6). ELISA assays were then performed to evaluate the ability of CoV-X2 to inhibit the binding of recombinant ACE2 to the S trimer (Fig.1e,f). In line with the structural information^10^, C135 did not affect the ACE2/S interaction. C121, which occupies the ACE2 binding site on the RBD, prevented ACE2 binding but only partially. By contrast, ACE2 binding was not detected in the presence of CoV-X2, suggesting a synergistic effect by the two moieties composing the bispecific.

To assess the neutralizing ability of CoV-X2 *in vitro*, we first used SARS-CoV-2 pseudoviruses^16^. The bispecific neutralized pseudovirus carrying wild-type SARS-CoV-2 S at sub-nanomolar concentrations (IC_50_ = 0.04 nM (5.8 ng/mL); IC_90_ = 0.3 nM (44 ng/mL)), which was similar or better than the parental IgGs and >100-fold better IC_50_ than the parental Fabs (Fig.1g). CoV-X2 remained effective against pseudoviruses bearing escape mutations that made them resistant to the individual mAbs (Fig.1g)^15^ and against a pseudovirus with RBD mutations found in the B.1.351 variant (first reported in South Africa, IC_50_ =1.3 nM (191 ng/mL); Extended data Fig. 5). To confirm CoV-X2 efficacy, we performed plaque reduction neutralization assays with infectious virus. CoV-X2 efficiently neutralized: SARS-CoV-2 (IC_50_ = 0.9 nM); the D614G variant first appearing in Europe (B.1, IC_50_ = 0.2 nM); the B.1.1.7 variant first observed in the United Kingdom (IC_50_ = 0.2 nM); the B.1.1.248 variant first isolated in Brazil (IC_50_ = 2.1 nM) and B.1.351 first isolated in South Africa (IC_50_ = 12 nM; Fig.1h). The latter two have almost identical mutations in the RBD, the only difference being N vs. T at position 417, which does not interact with CoV-X2. Nonetheless, neutralization of B.1.351 was lower, suggesting either some conformational differences in the RBD or long-range effects deriving from other mutations in the S protein. A similar behavior is seen with the wild-type sequence (D614), which has lower neutralization than G614 even if no other difference is present; a plausible explanation is that G614 makes the CoV-X2 epitopes more accessible by favoring the RBD ‘up’ conformation.^17^ We conclude that the *in vitro* binding and neutralizing properties of CoV-X2 make it preferable over its parental antibodies.

To assess the clinical potential of CoV-X2, we investigated its ability to protect animals from infection and disease. We first developed a novel mouse model in which human ACE2 (hACE2) is expressed by upper and lower respiratory tract cells upon inhalation of a modified Adeno Associated Virus (AAV-hACE2, see Methods, Fig.2 and Extended Data Fig.7).

**Fig. 2.**
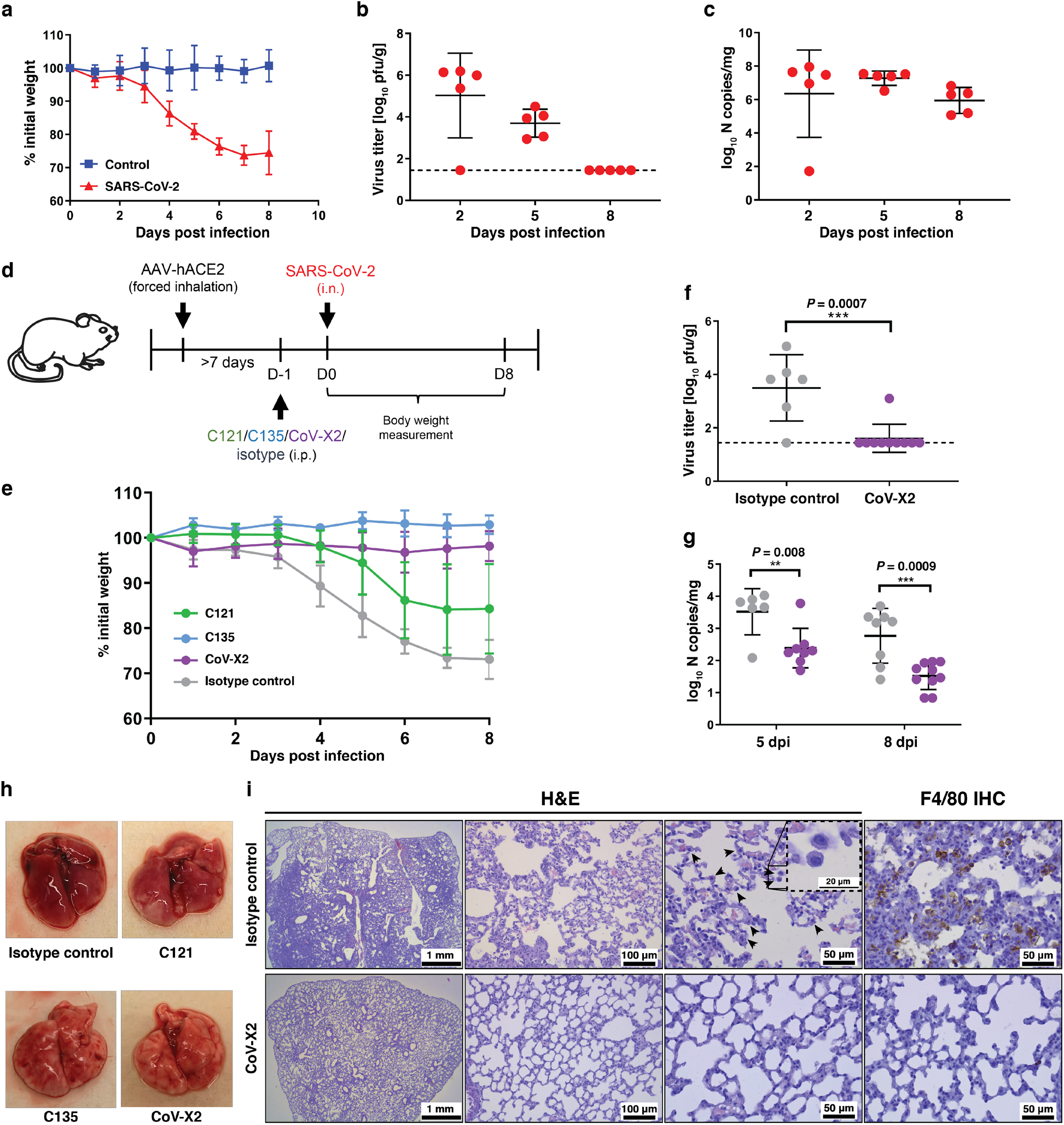
CoV-X2 protects AAV-hACE2-transduced mice against SARS-CoV-2 disease. **a**, Loss of body weight over time in SARS-CoV-2 infected mice. 13 to 15 weeks old C57Bl/6NCrl wild-type female mice were transduced with AAV-hACE2 by forced inhalation, which provides delivery of viral particles to both upper and lower respiratory tract. After >7 days, mice were either infected with SARS-CoV-2 (1×10^4^ pfu) or received vehiculum by the intranasal route. Weight was monitored daily for 8 days (SARS-CoV-2, n = 5; control, n = 4). Mean with standard deviation is shown. **b**, Kinetic of viral burden in the lungs from SARS-CoV-2-infected mice by plaque assays. Mean with standard deviation; the dashed line indicates the limit of detection. **c**, Kinetic of viral RNA levels in lung samples from SARS-CoV-2-infected mice by RT-qPCR. Mean with standard deviation. **d**, Schematic of the experimental layout. Wild-type mice were transduced with AAV-hACE2 by forced inhalation. After >7 days, mice were inoculated intraperitoneally (i.p) with 150 µg of antibodies. One day later, the mice were infected intranasally (i.n.) with SARS-CoV-2 (1×10^4^ pfu). **e**, Changes in body weight upon infection were monitored daily in antibody-treated mice (C121, n=9; C135, n=5; CoV-X2, n=13; isotype control, n=10). Mean with standard deviation is shown. **f**, Lung viral burden by plaque assay at 5 dpi (isotype control, n=6; CoV-X2, n=10). The dashed line indicates the limit of detection; mean with standard deviation. P value was calculated with two-tailed Student’s t test. **g**, Spleen viral RNA levels by RT-qPCR at 5 and 8 dpi (gray: isotype control; purple: CoV-X2). Mean with standard deviation. P value was calculated with two-tailed Student’s t test. **h**, Photographs of lungs collected from infected mice (8 dpi). **i**, Histopathology and F4/80 immunohistochemistry (IHC). Hematoxylin and Eosin-stained (H&E) sections of paraffin-embedded lungs from infected mice (8 dpi). Arrowheads point to foamy macrophages. F4/80 IHC shows abundant macrophage infiltration in lungs of mice treated with isotype control but not with CoV-X2.

This approach enables rapid production of large cohorts of animals and has the advantage of being applicable to wild-type and mutant mouse colonies, independently of age and gender. Moreover, since AAV vectors are only weakly immunogenic and cytotoxic, the system allows for prolonged expression of hACE2^18-21^ (Extended Data Fig.7). SARS-CoV-2 infection of ACE2 humanized mice results in progressive weight loss, respiratory pathology and disease requiring culling on day 8 post infection (dpi, Fig.2a–c and Extended Data Fig.7).

To evaluate the protective effect of antibodies, hACE2 mice were treated with antibody (150 µg) one day before SARS-CoV-2 challenge and monitored over time (Fig.2d–i). Upon intranasal infection with 1×10^4^ pfu of SARS-CoV-2 (SARS-CoV-2/human/Czech Republic/951/2020), isotype control treated animals showed weight loss starting at 3 dpi, and by 8 dpi most animals had lost approximately 25–30% of their body weight reaching humane endpoint (Fig.2e). Infectious virus could be recovered from the lungs (Fig.2f), viral RNA was detected also in the spleen (Fig.2g) but not in the heart (data not shown). Lung pathology resembled severe COVID-19 in humans^22^ and was characterized by Diffuse Alveolar Damage (DAD; 50-80% of tissue area), alveolar replacement with infiltrates of immune cells and fibroblasts, thickened septa and infiltrations by activated macrophages with foamy cytoplasm (Fig.2i). In contrast, animals treated with CoV-X2 maintained their body weight (P<0.0001 at 4–8 dpi when compared to isotype; Fig.2e; P values between all groups in Extended Data Table1), had reduced viral RNA in the spleen (Fig.2g) and displayed neither macro-nor histopathological changes (DAD <5-10%, Fig.2h,i). While infectious virus could be readily recovered from controls (5 of 6), it was only recovered from 1 out of 10 CoV-X2 treated animals at 5 dpi (Fig. 2f) and could not be recovered from any of 13 animals at 8 dpi (data not shown). Since none of the CoV-X2 treated mice exhibited symptoms at any time, we conclude that CoV-X2 protects mice from infection and disease.

Since monotherapy with C121 or C135 mAbs leads to virus escape *in vitro*^15^, we treated hACE2 mice with the individual antibodies and sequenced the virus. Only wild-type RBD sequences were obtained from controls (n=10). Instead, the virus in mice treated with C121 selected for a mutation resulting in E484D (5 of 5 mice that were analyzed at 8 dpi). C121 escape mutations at E484 were previously observed *in vitro*^15^ and changes at this residue (present also in the B.1.351 and B.1.1.248 variants) reduce neutralization by human sera by more than 10-fold^23^. E484D affects intermolecular H-bonds at the core of the C121/RBD interface and it is suggested to increase the RBD affinity for ACE2^24^. Virus with D484 is pathogenic, since 7 out of 9 mice treated with C121 developed disease (Fig.2e) and only D484 virus was found in their lungs. In contrast, and unlike the *in vitro* results^15^, no virus evasion or pathology was observed in mice treated with C135 (n=5; Fig.2e and data not shown). In CoV-X2 treated animals, even though no infectious virus was retrieved (8 dpi, n=13) and no symptoms ever noticed, low levels of residual viral RNA could be detected in some animals after 40 cycles of PCR amplification: in 6 of 13 animals the virus sequence was wild-type and in 2 mice overlapping sequencing traces were consistent with coexistence of wild-type and D484. Thus, in those 2 of 13 animals with D484 CoV-X2 remained protective even if the mutation diluted the effective antibody concentration, presumably leaving only the C135 moiety active. Finally, CoV-X2 was protective also when administered 12 hours after SARS-CoV-2 challenge (Extended Data Fig.8)

Monoclonal antibodies targeting the SARS-CoV-2 S are in advanced clinical trials and show promise against COVID-19^1,2^. Concomitant use of multiple antibodies is preferred for increased efficacy and added resistance against viral evasion. Indeed, the virus can escape pressure by a single antibody *in vitro* and, as shown here, also in animals. Moreover, RBD mutations threatening the efficacy of single monoclonals have already been detected in virus circulating in minks and humans^25^, including mutations at the C121 and C135 epitopes (Extended Data Fig.9). One disadvantage of antibody cocktails is the requirement for twice or more the development and production capacity than for single mAbs, which is a significant challenge in light of the augmented demand due to COVID-19 related vaccines and therapeutics on top of the need to maintain production of biologics for other diseases.^4^

Multispecific antibodies offer the advantages of cocktails in a single molecule. Indeed, we have shown that the CoV-X2 bispecific is more effective than the related monoclonals at inhibiting ACE2 binding; it has sub-nanomolar IC_50_ against a broader array of viral sequences; and it protects animals from SARS-CoV-2 even when C121, its potent parental mAb, fails due to the insurgence of viral escape. C135, the other parental mAb, did not generate escape in our animal experiment but readily generated them *in vitro*^15^. CoV-X2 is expected to be more resistant to viral escape compared to monoclonals. Indeed, we have shown that CoV-X2 binds and neutralizes mutants not recognized by its parental mAbs as well as variants of concern that recently emerged in United Kingdom^13^, South Africa^14^ and Brazil^26^.

CoV-X2, unlike other multispecifics^27^, is a fully human IgG-like molecule. As such, it has favorable developability and could be further engineered to alter effector functions. For example, the Fragment crystallizable (Fc) of CoV-X2 was already modified to modulate its interaction with Fc receptors and complement (LALA-PG mutations)^28^ without affecting its antigen-binding properties. The LALA modification prevents Antibody Dependent Enhancement (ADE) of flavivirus infection^29,30^ and it may be a desirable modification also in the context of SARS-CoV-2, since cellular and animal experiments with coronaviruses, including SARS-CoV^31-33^, support the possibility of ADE. Other modifications, like LS^28^ for increased half-life, are easily achievable. Finally, CoV-X2 is human-derived and produced in a format (CrossMab) already shown to be safe in clinical trials^34^, which further supports its developability. Thus, IgG-like bispecifics are worth adding to the arsenal employed to combat SARS-CoV-2 and its plausible future mutations.

## Acknowledgements

*Dedicated to the memory of the recently departed Prof. François Diederich*.

This work was supported by: the European Union’s Horizon 2020 research and innovation program under grant agreement No. 101015756, ATAC consortium (EC 101003650; D.F.R., L.V., Q.P.H., F.B., L.C.); SNF grant 31003A_182270 (L.V.); Lions Club Monteceneri (L.V.); George Mason University Fast Grant (D.F.R.); NIH grant P01-AI138398-S1 (M.C.N., P.J.B.); 2U19AI111825 (M.C.N., D.F.R.); the Caltech Merkin Institute for Translational Research and P50 AI150464 (P.J.B.); R37-AI64003 (P.D.B.); R01AI78788 (T.H.); P.D.B. and M.C.N. are Howard Hughes Medical Institute Investigators. The study was also supported by: the Czech Academy of Sciences and Czech Ministry of Agriculture (RVO 68378050; R.S.; RVO0518; D.R.); Czech Ministry of Education, Youth and Sports and the European Regional Development Fund (LM2018126; CZ.1.05/2.1.00/19.0395 and CZ.1.05/1.1.00/02.0109; R.S.; CZ.02.1.01/0.0/0.0/15_003/0000495; D.R.); Czech Science Foundation (20-14325S, D.R.); and by Ricerca Finalizzata from Ministry of Health, Italy (grants no. GR-2013-02358399; A.P.). We are grateful for the high-performance computing resources provided by CINECA, Dr. Sanzio Bassini, to Prof. Michael Hust, Dr. Federico Bertoglio and Elisa Restivo. We thank Vaclav Zatecka, Veronika Martinkova, and Linda Kutlikova for technical assistance.

## Author contributions

R.D.G, M.Pe., L.S., F.Mu., J.C.L., F.Ma, D.M., C.I., E.P., S.D.G., M.Pa., F.B., D.M., S.Gi., C.O.B, F.B., J.C.S, F.G, S.Ga, designed and carried out experiments and analyzed results, produced plasmids, antibodies and viral proteins. P.N., T.M., J.H., V.H, B.M., N.P., A.F., J.T., V.I., M.Pa., D.Z., P.B., I.B., P.S., D.R., performed animal experiments and analyzed the results. L.V, D.F.R., D.R., Q.P.H., A.P., L.C., P.J.B., M.C.N., P.D.B., T.H. conceived and designed study and experiments and analyzed the results. P.N., T.M., R.N., O.P., J.P., J.R., R.S. conceived and designed the mouse model. L.V., D.F.R., D.R, R.D.G. wrote the manuscript with input from all co-authors.

## Extended data figures

**Extended Data Fig.1.**
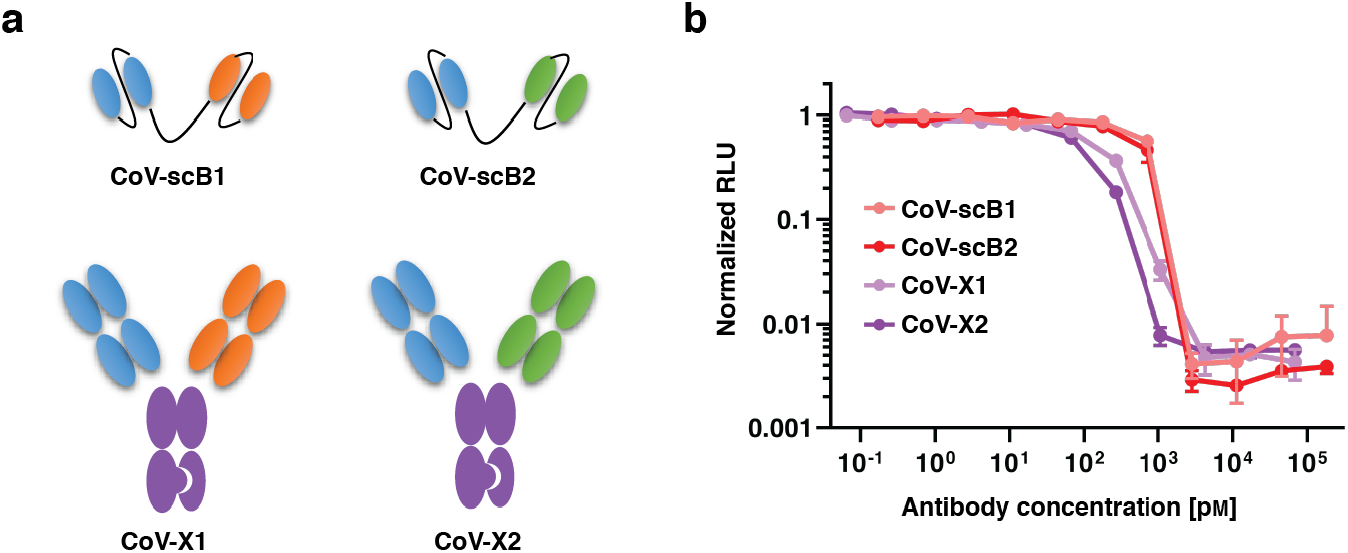
Neutralization of SARS-CoV-2 pseudovirus by bispecific antibodies. **a**, Schematic representation of the 4 bispecific constructs; two in scFv format and two as IgG-like CrossMAb with knob-in-hole. The parental monoclonals forming the bispecifics are color-coded (C135 blue, C144 orange, C121 green; Fc region in purple). **b**, All 4 constructs neutralize SARS-CoV-2 pseudovirus *in vitro* at sub-nanomolar concentrations (IC_50_: 0.13, 0.04, 0.74 and 0.53 nM for CoV-X1, CoV-X2, CoV-scB1 and CoV-scB2, respectively). Normalized relative luminescence values, which correlate to infection, are reported versus antibody concentration, as detailed in Schmidt *et al*.^16^. Mean with standard deviation is shown, representative of two independent experiments.

**Extended Data Fig.2.**
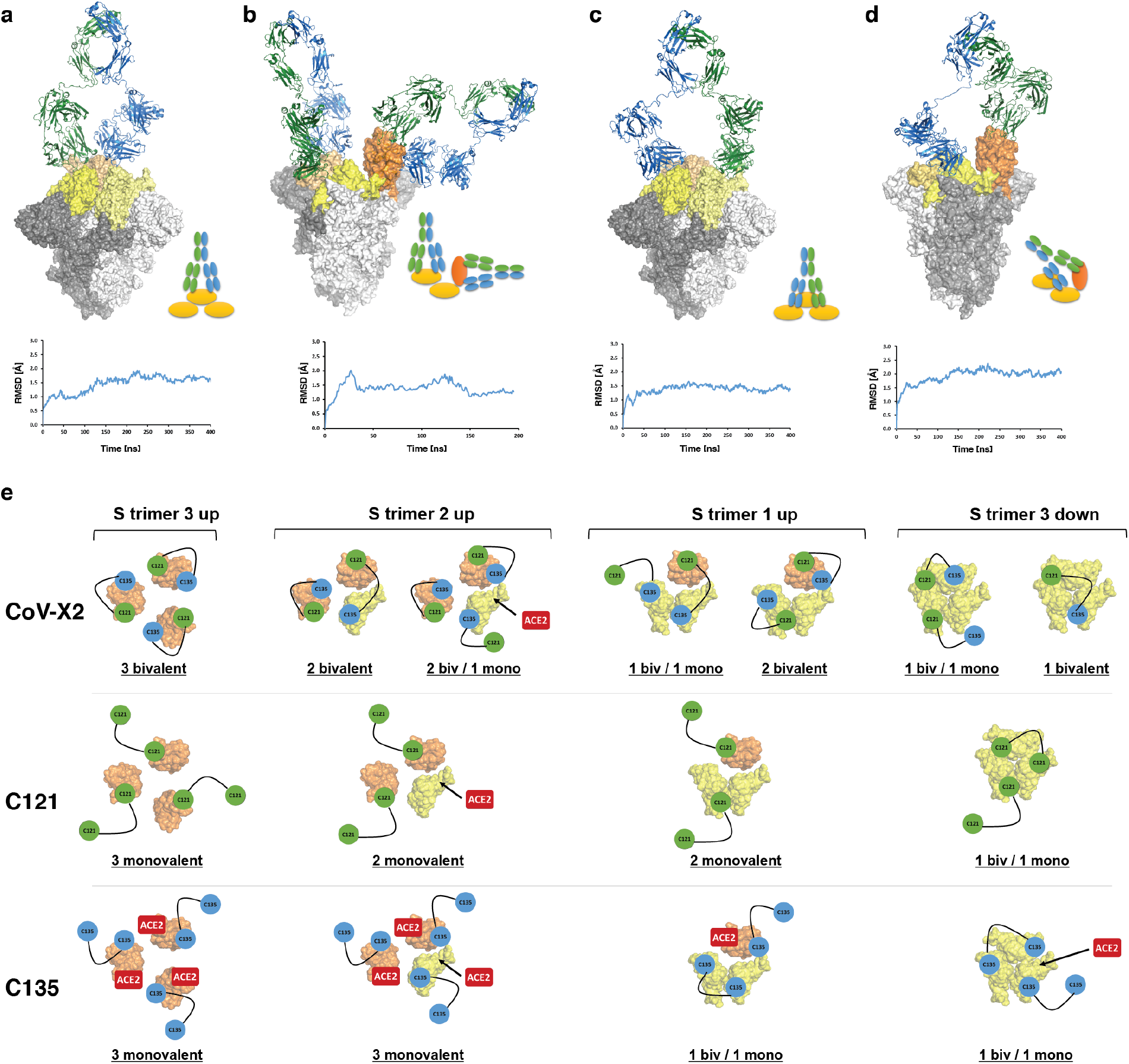
CoV-X2 engages its epitopes on all RBD conformations on the S trimer. **a–d**, Molecular Dynamics (MD) simulations of the complex between the CoV-X2 bispecific and S trimers with RBD in either all down, all up or mixed up/down conformations show that CoV-X2 can engage a single RBD with both arms (**a**,**b**), two adjacent RBDs in the down conformation (**c**), and two RBDs in the up/down conformation (**b**,**d**). The complexes were subjected to up to 400 ns of fully atomistic MD simulations to assess feasibility and stability of the bound conformations. Root-mean-squared deviations (RMSD) values are shown to indicate structural stability. S trimer is in shades of grey, RBDs in yellow (down conformation) and orange (up), the C121 and C135 moieties of CoV-X2 are in green and blue, respectively. **e**, Schematic representation of the computationally predicted binding modes of CoV-X2, C121 IgG and C135 IgG on the S trimer, colored as in **a**–**d**. Antibodies are represented by connected circles; ACE2 is in red on the RBD if it can bind directly to a given conformation; it has an arrow pointing to the RBD if ACE2 binding is achieved after an allowed switch to the up conformation. For example, in the 3-up conformation (left), CoV-X2 can engage all the RBDs with bivalent binding, whereas C121 and C135 can only achieve monovalent binding. C135 binding does not prevent interaction with ACE2. The situation is similar in the other S conformations (2-up 1-down, 2-down 1-up and 3-down), with only the bispecific achieving bivalent interaction and preventing ACE2 access in all conformations.

**Extended Data Fig.3.**
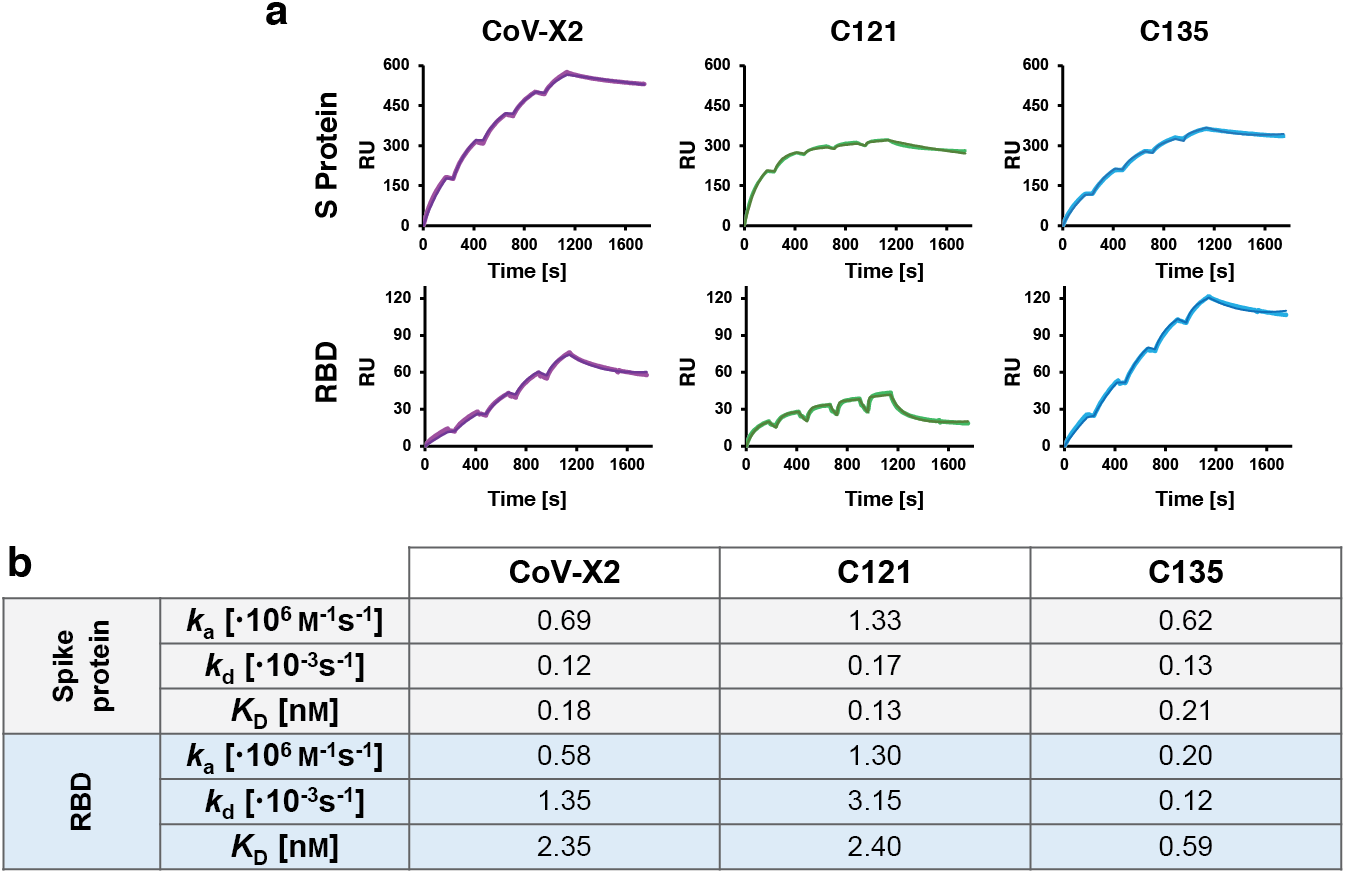
CoV-X2 and its parental mAbs bind recombinant, isolated RBD and S trimer with low nanomolar affinity. **a**, Representative SPR traces from which the data in (**b**) was derived. **b**, Kinetic parameters for the binding of C121 IgG, C135 IgG, and CoV-X2 to S trimer and RBD.

**Extended Data Fig.4.**
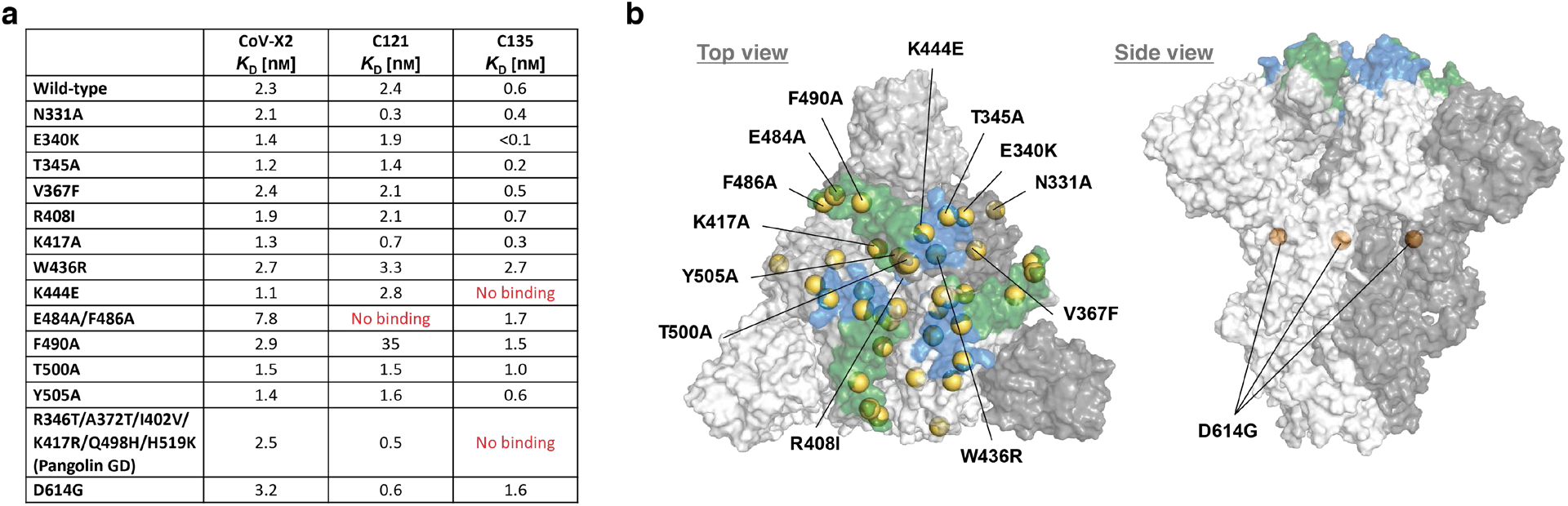
CoV-X2 binds with low-nanomolar affinity to S protein mutants, including some that are not recognized by the parental mAbs C121 and C135. **a**, SPR-derived binding affinities of CoV-X2, C121 IgG and C135 IgG to several S trimer mutants. **b**, Mutations tested in (**a**) are indicated by yellow spheres on the surface representation of the S trimer. The epitopes of C121 (green) and C135 (blue) are shown.

**Extended Data Fig.5.**
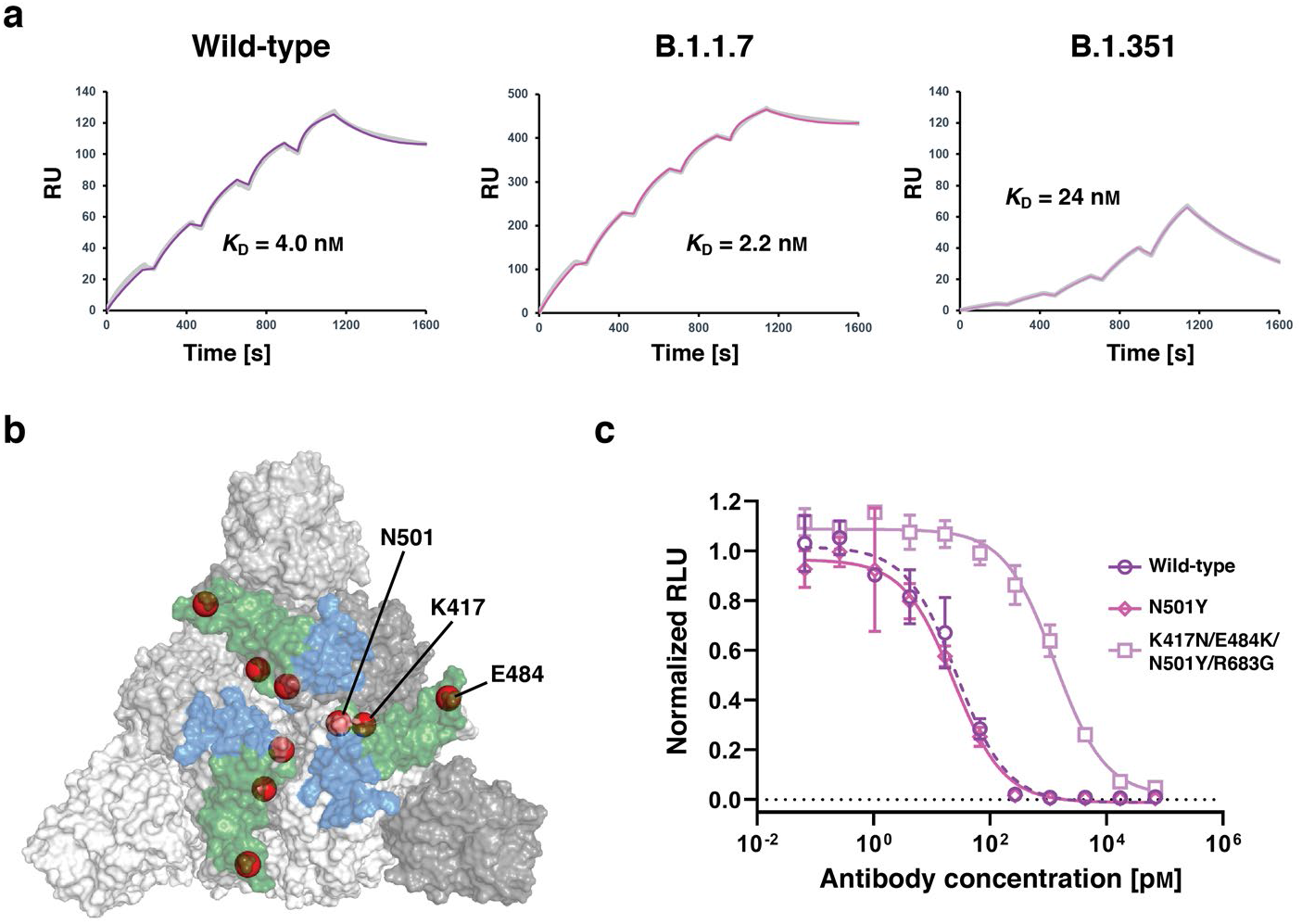
Efficacy of CoV-X2 against B.1.1.7 and B.1.351 variants. **a**, SPR traces showing binding of CoV-X2 to the RBD corresponding to wild-type, B.1.1.7 (also known as UK) and B.1.351 (also known as South African) variants of SARS-CoV-2. **b**, Residues mutated in the variants are shown as red spheres on the surface representation of the S trimer. The epitopes of C121 (green) and C135 (blue) are shown. **c**, Neutralization of SARS-CoV-2 pseudoviruses expressing wild-type, N501Y and K417N/E484K/N501Y/R683G (corresponding to South African mutants in the RBD, see Figure 1h) S protein by CoV-X2.

**Extended Data Fig.6.**
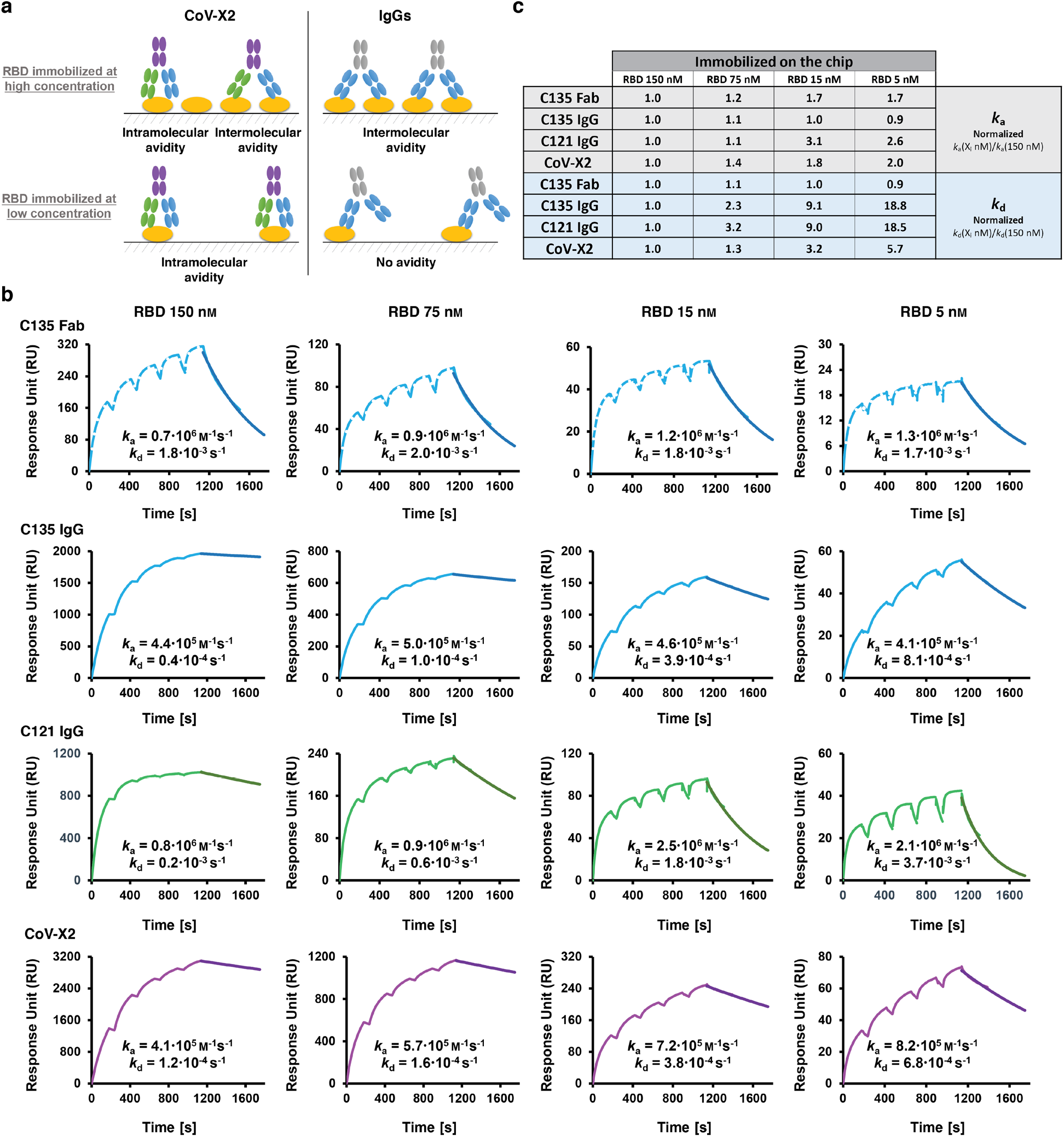
SPR-based avidity assays confirm that CoV-X2 can engage bivalently on a single RBD. **a**, CoV-X2 and monoclonal IgGs (C121 or C135) have different binding modes available when high or low quantities of RBD are immobilized on the surface of the SPR chip. mAbs have avidity effects at high RBD concentrations due to intermolecular binding, which results in slower dissociation rate (*k*_d_), but not at low RBD concentrations, since bivalent binding to a single RBD is impossible. In contrast, the bispecific has avidity at both high and low concentrations, since bivalent binding to its two epitopes on a single RBD is possible. *k*_a_ is not affected by avidity. **b**, Experimental confirmation that CoV-X2 engages bivalently on a single RBD. SPR traces used to determine *k*_a_ and *k*_d_ of mAbs, Fab and bispecific at different concentrations of immobilized RBD (see Fig.1d) are shown. **c**, Table summarizing the SPR results plotted in Fig.1d. *k*_a_ and *k*_d_ were normalized against the values at the highest RBD concentration. *k*_a_ and Fab *k*_d_ were unaffected by the RBD concentration, as expected. *k*_d_ became faster for the monoclonals (loss of avidity) but less so for the bispecific (avidity maintained due to simultaneous binding to two sites on a single RBD).

**Extended Data Fig.7.**
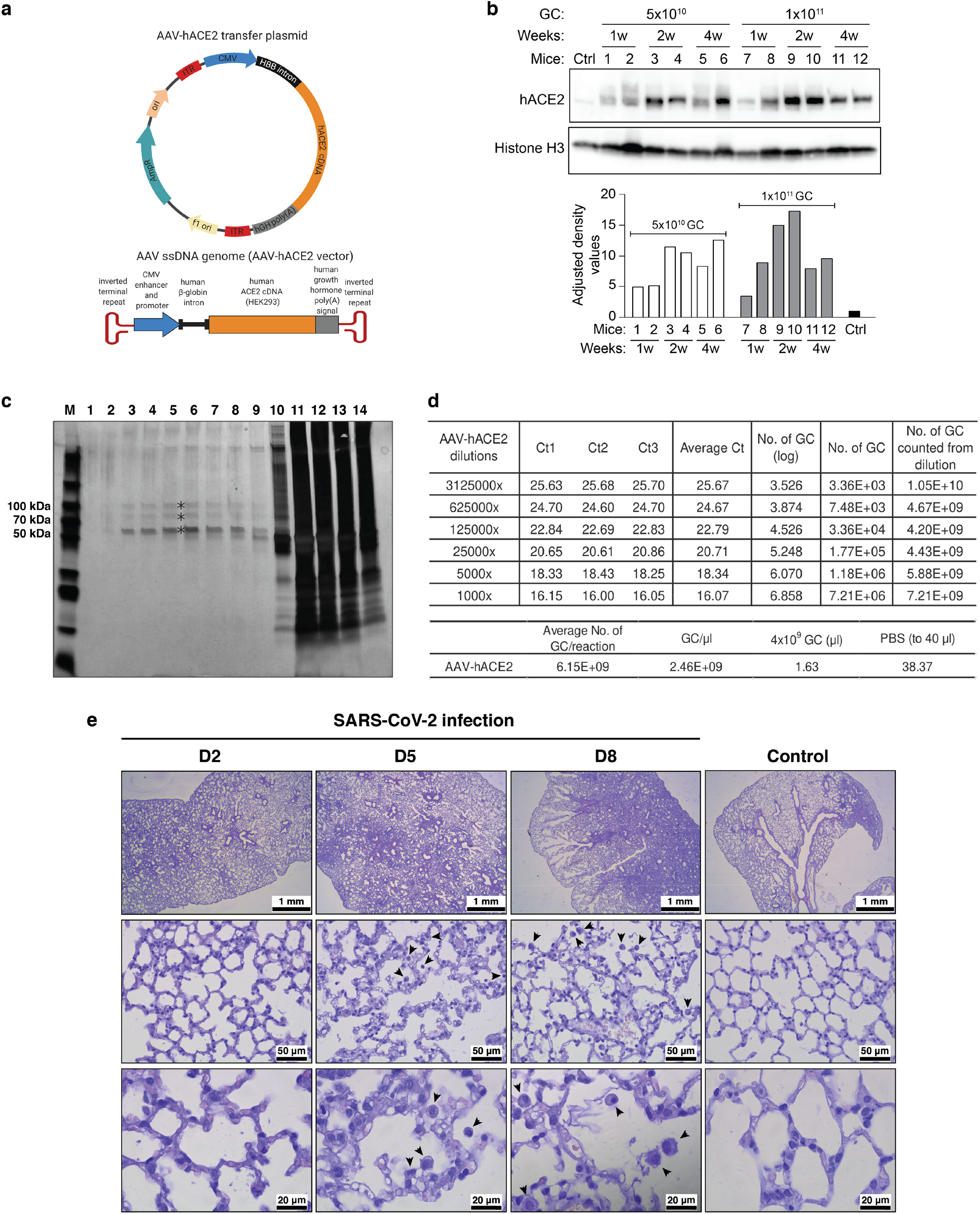
Generation of the new AAV-hACE2-transduced mouse model for COVID-19. **a**, Diagram of the AAV-hACE2 plasmid and corresponding Adeno Associated viral vector. **b**, Western blot analysis detecting hACE2 expression in the lungs of one non-transduced control mouse (Ctrl) and 12 mice transduced with two different doses of AAV-hACE2 viral particles (5×10^10^ or 1×10^11^ genome copies (GC)). Lung tissue was collected 1, 2, or 4 weeks (w) post transduction. Histone H3 was used as control for quantification (bottom). **c**, Preparation of concentrated AAV-hACE2. AAV-hACE2 plasmid was co-transfected with pHelper and AAV Rep/Cap 2/9n vectors into 293AAV cells (see Methods). In order to increase viral titers, viral particles from both cell lysate and PEG-precipitated growth medium were ultracentrifuged in discontinuous iodixanol gradient. The silver-stained SDS-PAGE gel shows 14 consecutive fractions: 1-9 represent enriched AAV fractions used for experiments, whereas fractions 10– 14 are contaminated with proteinaceous cell debris. Iodixanol was chosen as a density gradient medium due to its low toxicity *in vivo* and its easy removal by ultrafiltration. M is protein marker, * are AAV capsid proteins VP1, VP2, and VP3. **d**, The amount of AAV particles was estimated by qRT-PCR. The number of genome copies (GC) expressed as log was calculated from a standard curve. From one 15 cm^2^ dish, 75 µl with 2.0×10^12^ GC/ml were prepared, which is sufficient for hACE2 humanization of 37 mice. **e**, Kinetic of lung histopathology in SARS-CoV-2 infected ACE2 humanized mice. Hematoxylin and Eosin-stained sections showed inflammatory infiltrates composed of lymphocytes, macrophages, neutrophils, and fibroblasts replacing the alveoli. The size of the affected areas increased over time (area of diffuse alveolar damage: control <5-10%, 2 dpi <10-30%, 5 dpi 20-80 %, 8 dpi 50-90%). Alveolar septa were thickened in areas close to infiltrates. In samples collected at 5 and 8 dpi, an increased number of activated macrophages with foamy cytoplasm (black arrowheads) was seen. AAV-hACE2 transduced, SARS-CoV-2 uninfected mice were used as control and showed no significant pathology.

**Extended Data Fig.8.**
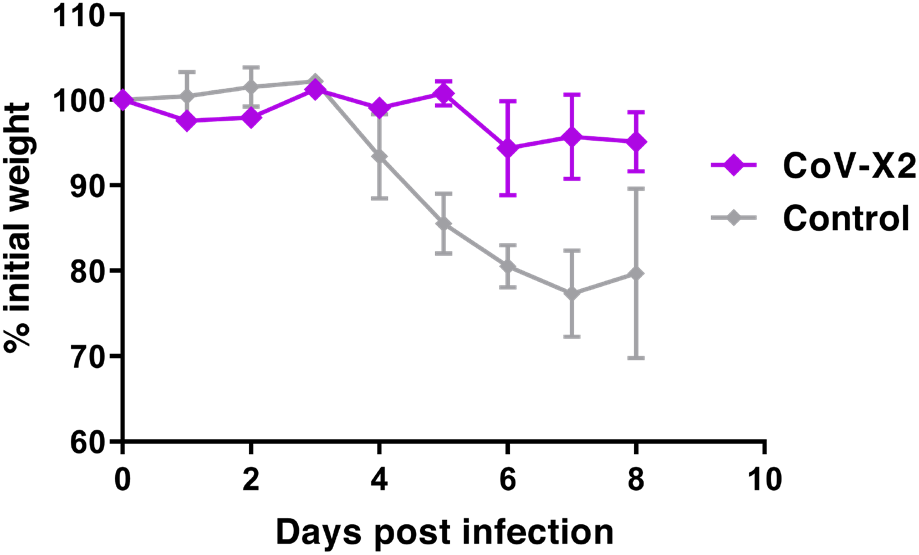
Post-exposure administration of CoV-X2 protects SARS-CoV-2 infected mice from disease. Animals were infected intranasally with 10^4^ pfu of SARS-CoV-2 and treated with 250 µg/mouse of either isotype control antibody (n=3) or CoV-X2 (n=2) 12 hours later. Weight loss and pathological signs were apparent in control but not in CoV-X2 treated animals.

**Extended Data Fig.9.**
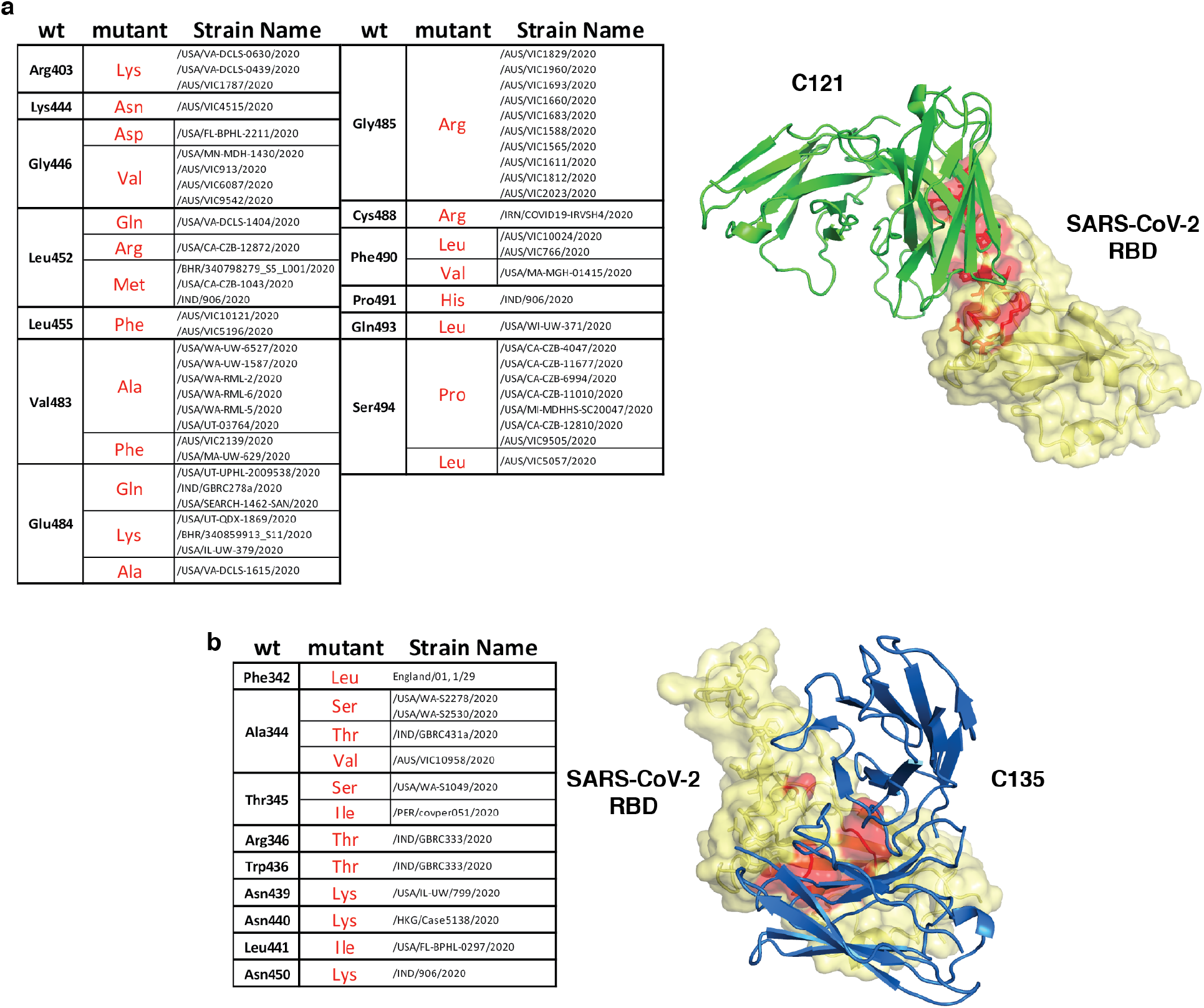
Natural SARS-CoV-2 variants in the C121 and C135 epitopes. Summary of naturally occurring mutations in the C121 (**a**) or C135 (**b**) epitopes reported in circulating SARS-CoV-2 (as of January 1, 2021). The location of the mutated residues is shown in red on the RBD structure. C121 and C135 variable regions are in green and blue (PDB ID: 7K8X and 7K8Z respectively).

**Extended Data Table 1.** Summary of the P values for the mouse protection experiment. Statistical comparison of body weight differences in animals treated with the individual monoclonal antibodies (C121 or C135), the CoV-X2 bispecific or isotype control at 8 dpi (related to Fig. 2e). P values were determined with the ANOVA test. Comparison of the entire curves (Fig. 2e) by the One Sample Wilcoxon Test or by the ANOVA followed by Turkey-Kramer post-test reveals that the isotype control treated group is statistically different from any of the other groups (CoV-X2, C135, or C121; P<0.05).

|  | C121 | C135 | CoV-X2 | Isotype control |
| --- | --- | --- | --- | --- |
| C121 | — | P<0.0001 | P<0.0001 | P<0.01 |
| C135 | P<0.0001 | — | P>0.05 | P<0.0001 |
| CoV-X2 | P<0.0001 | P>0.05 | — | P<0.0001 |
| Isotype control | P<0.01 | P<0.0001 | P<0.0001 | — |

